# Comparative analysis of PDZ-binding motifs in the diacylglycerol kinase family

**DOI:** 10.1101/2023.06.15.545061

**Authors:** Boglarka Zambo, Gergo Gogl, Bastien Morlet, Pascal Eberling, Luc Negroni, Hervé Moine, Gilles Travé

**Author notes:** Co-corresponding authors: Boglarka Zambo, Hervé Moine, Gilles Travé.

## Abstract

Diacylglycerol kinases (DGKs) control local and temporal amounts of diacylglycerol (DAG) and phosphatidic acid (PA) by converting DAG to PA through phosphorylation in cells. Certain DGK enzymes possess C-terminal sequences that encode potential PDZ-binding motifs (PBMs), which could be involved in their recruitment into supramolecular signaling complexes. In this study, we used two different interactomic approaches, quantitative native holdup (nHU) and qualitative affinity purification (AP), both coupled to mass spectrometry (MS) to investigate the PDZ partners associated with the potential PBMs of DGKs. Complementing these results with site-specific affinity interactomic data measured on isolated PDZ domain fragments and PBM motifs, as well as evolutionary conservation analysis of the PBMs of DGKs, we explored functional differences within different DGK groups. All our results indicate that putative PBM sequences of type II enzymes are likely to be nonfunctional. In contrast, type IV enzymes possess highly promiscuous PBMs that interact with a set of PDZ proteins with very similar affinity interactomes. The combination of various interactomic assays and evolutionary analyses provides a useful strategy for identifying functional domains and motifs within diverse enzyme families.

## Introduction

Diacylglycerol kinases (DGKs) are evolutionary conserved lipid kinases whose enzymatic function is to convert DAG to PA by phosphorylation. In cells, they participate in lipid turnover and regulate the spatio-temporal amounts of DAG and PA. Both lipids act as secondary messenger molecules and play essential roles in mediating cell responses by recruiting and modulating the activity of DAG and PA-binding proteins *(1)*. All DGK family members are highly expressed in the brain and impaired functions of certain DGKs lead to neurological diseases such as epilepsy, obsessive compulsive disorder, schizophrenia, bipolar disorder or fragile X syndrome *(2–8)*. They are therefore promising targets for new therapeutic approaches in these diseases and understanding their regulation is of great interest *(9–11)*.

Mammals comprise ten DGKs classified into five types (types I-V), with a diverse variety of highly conserved domains and motifs within each type *(12)*. All DGK enzymes contain a catalytic domain (DAGK) composed of a catalytic (DAGKc) and an accessory (DAGKa) subdomains, that is responsible for the phosphorylation activity of the protein and at least two C1 domains that bind phorbol esters and DAG *(13)*. In addition, there are a large variety of modular domains and functional motifs that are unique to certain types of DGKs *(13)*. These protein regions serve specific functions, such as activity regulation and subcellular localization of the different members of the family *(12)*. For example, five out of the ten human DGKs contain putative PDZ-binding motifs (PBMs).

PDZ domains (named after three proteins that contain this domain, Postsynaptic density protein PSD-95, Drosophila disk large tumor suppressor DLG1, and Zonula occludens-1 protein ZO-1) can interact with PBMs generally located at the C-terminus of their partner proteins. PDZ domain-containing proteins are thought to be involved in the organization and localization of supramolecular signaling complexes in the cells *(14)*. Among DGKs, type II (DGKδ, η and κ) and type IV (DGK ζ and ι) carry a putative C-terminal tail segment that conforms to the class I (DGKη, κ, ζ, ι) or class II (DGKδ) PBM consensus sequences (class I: [S/T-X-Φ-$], class II: [Φ-X-Φ-$], class III: [E/D-X-Φ-$] where Φ is any hydrophobic amino acid, X is any amino acid and $ denotes the free COOH terminus, according to the Eukaryotic Linear Motif database *(15)*). Although PDZ-PBM interactions may be important in the regulation of all PBM-containing DGKs, only some PDZ-mediated interactions of the DGKζ and ι enzymes have been characterized in depth so far.

It has been shown that the PBM of DGKζ binds to the PDZ domains of syntrophins and together they form a ternary complex with dystrophin in cell bodies and dendrites in neurons *(16)*. Furthermore, DGKζ, syntrophin and the small GTPase Rac1 can form a signaling complex that mediates neurite outgrowth in neurons *(17)*. In skeletal muscle, syntrophins are important in the recruitment of DGKζ to filamentous actin and Rac at lamellipodia and ruffles and the alteration of this interaction may contribute to disease pathogenesis in dystrophic muscle *(18)*. DGKζ can also interact with all members of the multi-PDZ domain containing Discs-large (DLG) proteins and the interaction with the postsynaptic density protein 95 (PSD-95/DLG4) is required for spine maintenance *(19)*. Interestingly, it has been also shown that the other member of the type IV DGKs, DGKι can also interact with PSD-95/DLG4, but it has been proposed that DGKι acts rather presynaptically and regulates the presynaptic release probability during metabolic glutamate receptor-mediated long-term depression (mGluR-LTD) *(20)*. DGKζ also binds by a PDZ-mediated manner to Sorting Nexin 27 (SNX27), whose function is the fast recycling of plasma membrane proteins from endosomes, and this interaction has been shown to participate in vesicle trafficking, antigen induced transcriptional activation and metabolic reprogramming in T-cells *(21, 22)*. These data suggest that the PBMs of DGKs play a crucial role in the precise localization of the proteins in supramolecular complexes at specific subcellular regions, such as synaptic membranes, where they modulate receptor signaling events by converting DAG to PA locally, but we lack a comprehensive view of the PDZ binding repertoire of all PBM containing DGKs, which would be essential to elucidate their specific functions.

In this study, we comparatively analyze the functionality of the PBMs within the DGK family by using various interactomic approaches and evolutional conservation analysis. Our results reveal that, despite conforming to the consensus, the PBMs of type II enzymes fail to bind PDZ domains, whereas the PBMs of the type IV kinases exhibit a remarkably promiscuous binding profile and show close functional similarity.

## Results

### Quantifying apparent equilibrium binding constants from cell extracts by nHU-MS

To screen for new partners and simultaneously quantify dissociation constants across the proteome, we used our novel nHU-MS approach, where a PBM bait is probed against a whole cell lysate containing thousands of distinct preys *(23)*. As baits, we used biotinylated 10-mer C-terminal PBM peptides of the five type II and type IV DGK enzymes (Fig. 1A-F) or biotin as control, both immobilized at 10 μM final concentration, and measured the depletion ratio of full-length partners directly from neuroblastoma SH-SY5Y cell lysates after two hours of incubation to ensure binding equilibrium. The measured relative prey concentration difference in the liquid phases of sample and control experiments can be converted to apparent affinities as we previously demonstrated *(23)*. In the experiment series, 4,148 full-length endogenous proteins were detected and a total of 10,429 affinity measurements were performed with the five PBMs (Table S1). Hyperbolic binding thresholds were applied to separate significantly depleted proteins from the background, considering both the enrichment values and the statistical variance of the measurements *(24)*. Significant binding, albeit with low affinity and statistical confidence, was found for DGKδ, DGKκ and DGKη in 15, 14 and 19 partners, respectively, among which no PDZ domain-containing protein was found (Fig. 1A-C). For DGKζ and DGKι, we have detected 23 and 13 significant binders, out of which 5 were found to be PDZ domain-containing proteins (Fig. 1D-E). These 5, namely SNX27, MAGI3, LIN7C, CASK and SYNJ2BP, were all shared partners of DGKζ and DGKι. By correlating the measured affinities for DGKζ and DGKι, which were statistically significant for DGKζ, we found a significant correlation between the affinities of the two PBMs, suggesting that the two PBMs are functionally closely related (Fig. 1F).

**Figure 1.**
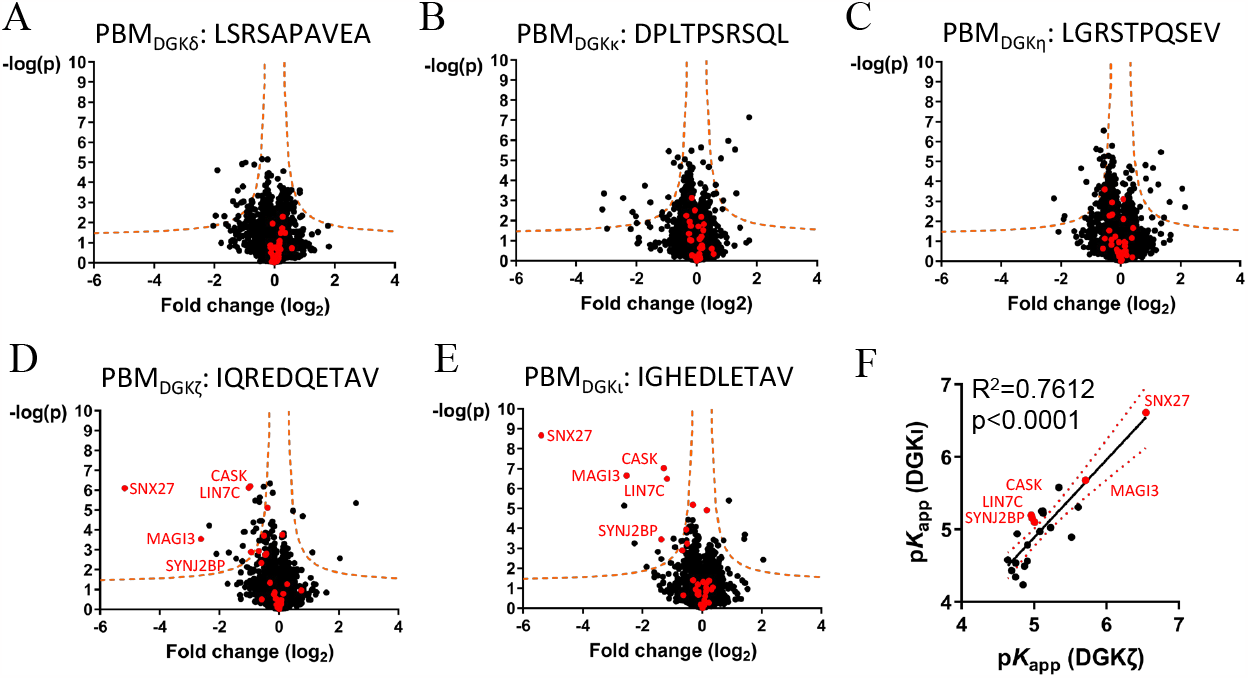
Binding partners of the C-terminal PBMs of the different type II (δ, κ and η) and type IV (ζ and ι) DGKs determined by nHU-MS on SH-SY5Y cell lysates. **A-C.** Volcano plots of nHU-MS with the PBM of type II DGKs, DGKδ (A), DGKκ (B) and DGKη (C). **D-E**. Volcano plots of nHU-MS with the PBM of type IV DGKζ (D) and DGKι (E). In the case of binding, a depletion is expected in the volcano plots, so the binders appear on the left side of the plot. PDZ domain-containing proteins are labelled red on all plots. Orange dashed curves: significance thresholds. **F**. Correlation between the apparent binding affinities measured for DGKζ and DGKι peptides. Red dotted line: 95% confidence interval. R^2^: Pearson correlation coefficient, p: p-value of two-tailed t-test, p*K*_app_ = −log_10_(*K*_app_ [M]).

### Exploring additional interaction partners from cell extract by AP-MS

We proved before that nHU is a sensitive method to detect and quantify even weak and transient interactions *(23)*. However, as there is no enrichment of partners in this assay it can miss important interaction partners if they have low abundance, *i*.*e*. below the detection threshold of total proteomic measurement. For this reason, we performed pulldown-based AP-MS experiments to verify that type II enzymes do not bind any PDZ proteins from the proteome, while type IVs are promiscuous binders. We used the biotinylated PBMs of DGKκ and DGKζ from the two groups and used SH-SY5Y cell lysates as before (Fig. 2A-B). The relative amounts of 713 proteins were determined in the experiment series (Table S2). We identified 15 prey proteins for DGKζ, and only 1 for DGKκ peptide bait in the AP-MS experiments above the hyperbolic significance threshold (Fig. 2A-B, Table S2). The sole potential partner detected for DGKκ peptide bait (TOMM22) does not contain any PDZ domain and is also enriched in the DGKζ experiment indicating that it is most likely false positive. Overall, we detected 22 PDZ domain-containing proteins in the experiment series, of which 12 showed significant enrichment on the DGKζ bait saturated resin. Among the PDZ partners identified by nHU-MS of DGKζ, only CASK was not recognized as a significant partner in AP-MS. In addition to the PDZ domain, CASK contains heterotetramerization L27 domains, which are exclusively found in PDZ proteins leading to co-enrichment of multi-component PDZ complexes, which may explain its binding in nHU-MS *(25, 26)*. Furthermore, MMP3 and MMP5 have been found to bind according to AP-MS, but not in nHU-MS. These proteins also possess L27 domains, which could potentially contribute to their indirect co-purification through stronger partner proteins that also contain L27 domains, such as DLG1.

**Figure 2.**
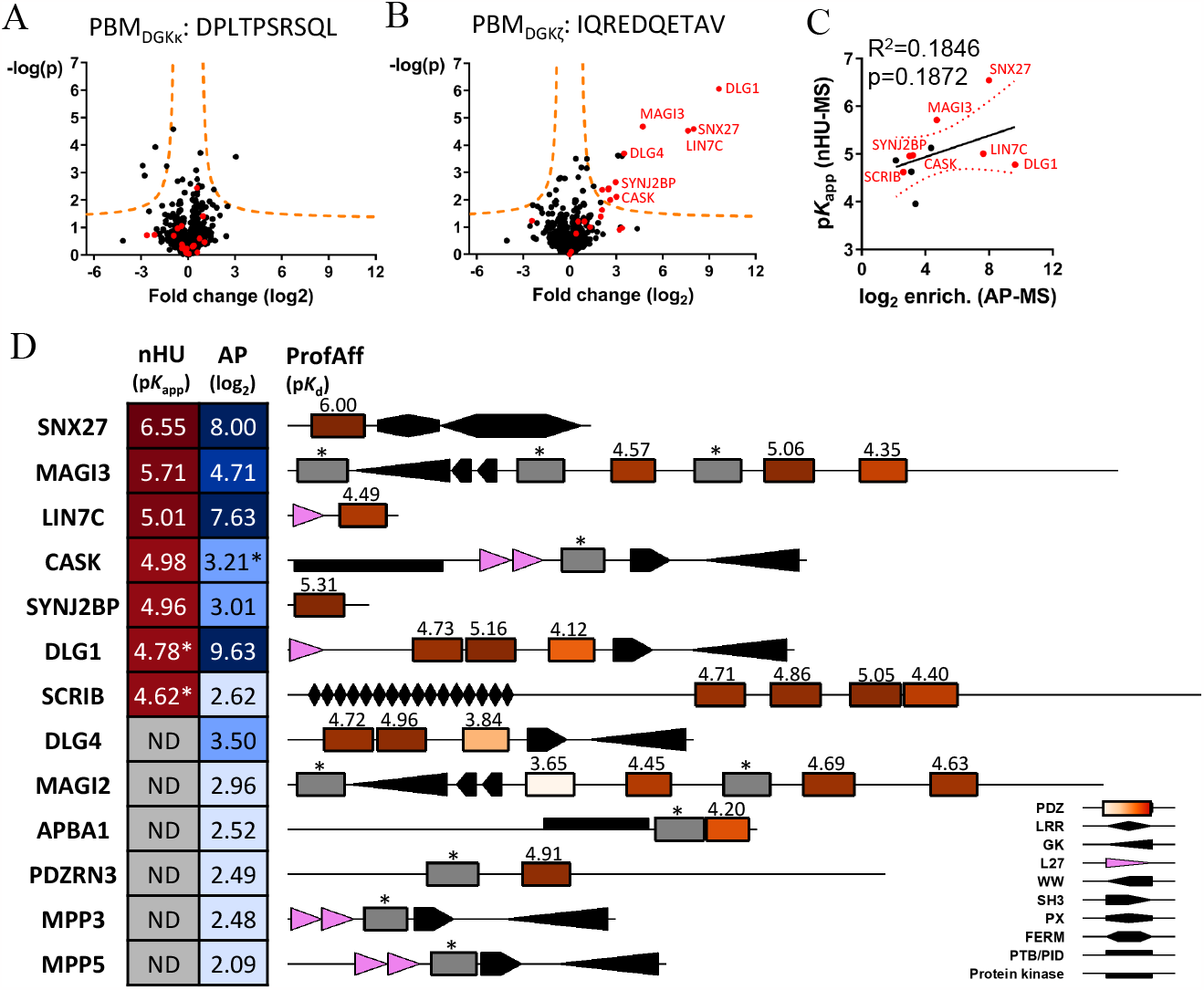
Binding partners of the C-terminal PBMs of DGKκ (type II) and DGKζ (type IV) determined by AP-MS on SH-SY5Y cell lysates. **A.** Partners detected for the DGKκ peptide. Only one partner seems significant according to our thresholds, and it does not contain any PDZ domain. **B**. Partners detected for the DGKζ peptide. 15 significant binding partners were detected, among which 12 contain PDZ domains. **C**. Comparision of measured enrichment values in AP-MS and the determined p*K*_app_s in nHU-MS for the proteins detected by both methods. PDZ domain-containing proteins labelled red on panels A-C. **D**. The domain structures of major PDZ domain-containing interactors of the DGKζ PBM according to the two interactomic assays (nHU-MS, AP-MS) that has been performed in this study. Color codes from yellow to red indicates weaker to stronger affinity values (nHU, ProfAff) or from light blue to dark blue weaker or higher enrichment values (AP). Grey: Not detected (ND) or below threshold (*). Values over PDZ domains: p*K*_d_s from ProfAff database^25^. Visualization: ProFeatMap^28^.

Compared to nHU-MS experiment, there are more PDZ domain-containing proteins among significant binders in the AP-MS with the DGKζ peptide (5/23 in nHU and 12/15 in AP, Table S3). Between the enrichment values of proteins detected in AP-MS and the determined p*K*_d_ values by nHU-MS, we could not find a correlation (p*K*_d_ =□log_10_(*K*_d_ [M]), Fig. 2C). Given that the two methods measure different parameters of interactions, this result is not surprising *(27)*. Overall, by combining the two different techniques, we demonstrated, that type II enzymes are apparently unable to recruit PDZ-domains and we have determined the affinities and validated the key PDZ-mediated interactions of type IV enzymes in neuroblastoma cells (Fig. 2D, visualization by ProFeatMap *(28)*, site-specific affinities by ProfAff database *(25)*).

### Correlative analysis of fragmentomic PDZ-binding profiles of different DGKs using the ProfAff database

Both nHU-MS and AP-MS measure interactions with full-length proteins across the proteome, yet the utilized motifs are expected to bind only with PDZ domain-containing proteins in a direct manner. To compare the binding profiles of the PBMs of DGKs across the entire human PDZome in the most sensitive manner, we analyzed the recently measured PDZ-PBM affinity interactomic map (available on the ProfAff server, *(25)*). These data consist of equilibrium binding constants of >65,000 PDZ-PBM interactions that were measured between purified PDZ domains and synthetic 10-mer C-terminal PBM peptides by high-throughput holdup method *(25)*. The binding constants for DGKκ (type II) and DGKζ (type IV) were determined against 97% of all the PDZ domains of the human proteome (258 out of 266, Table S4). For the DGKζ peptide, 26% of the total measured PDZome showed binding above the detection threshold and several strong binding affinities were determined (13 PDZ domains showed p*K*_*d*_ > 5), while for the DGKκ peptide only 9% of the measured PDZome showed detectable binding and all interactions determined were below 5 p*K*_*d*_ (Fig. 3A). We have found only minor overlap between the identified binders (Jaccard similarity=0.18) and no significant correlation between the affinities of the binders for DGKζ and DGKκ (n=14 over 258, Pearson correlation coefficient R^2^=0.015, two-tailed t-test p=0.67, Fig. 3B).

**Figure 3.**
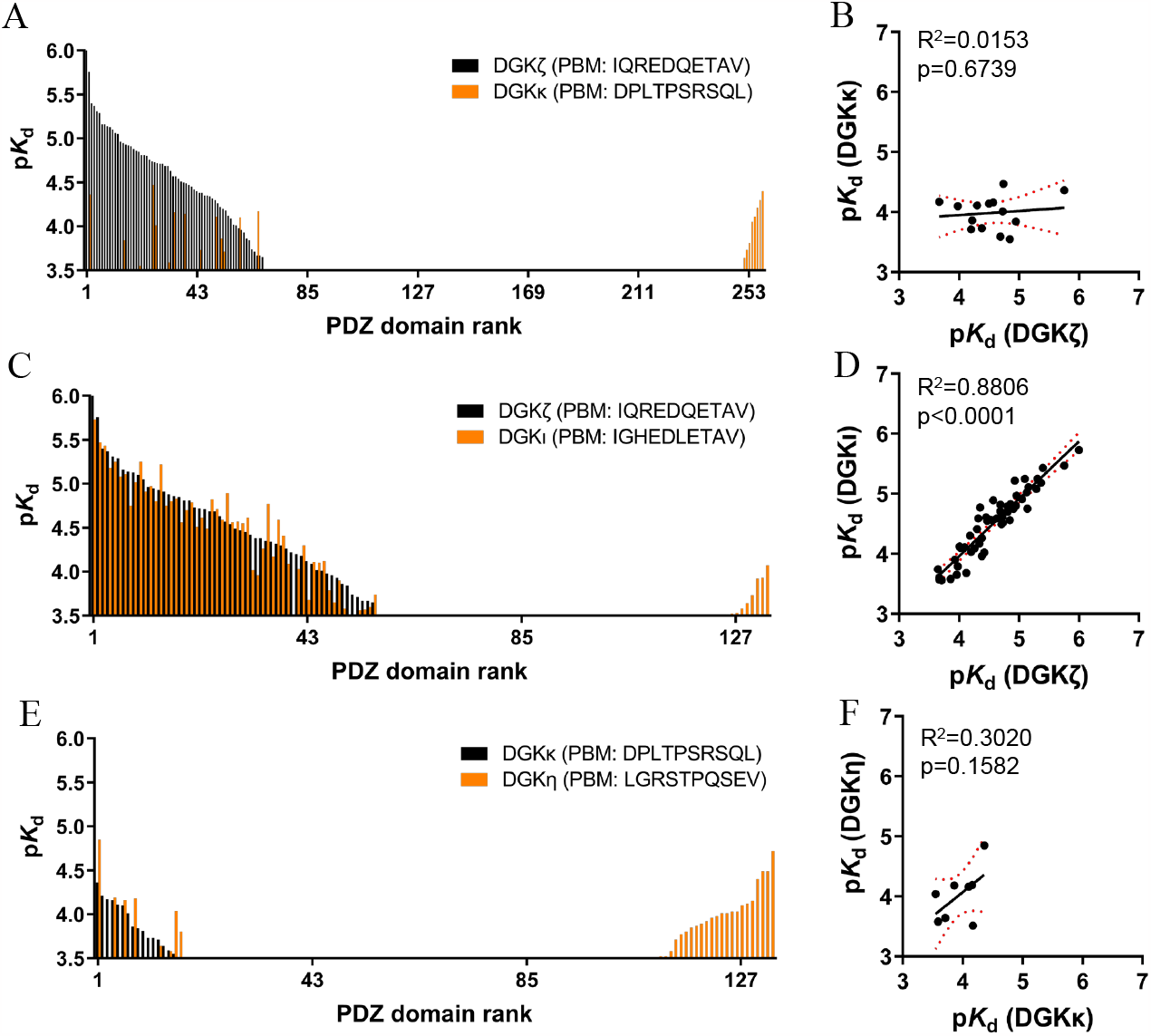
Comparative analysis of PDZ domain-binding profiles of PBM-containing DGKs using the ProfAff database. **A.** p*K*_d_ values measured between the PBMs of DGKζ (black) and κ (orange) and different PDZ domains. **B**. Correlation between the binding affinities measured for DGKζ and κ peptides. **C**. p*K*_d_ values measured between the PBMs of DGKζ (black) and ι (orange) and different PDZ domains. **D**. Correlation between the binding affinities measured for DGKζ and ι peptides. **E**. p*K*_d_ values measured between the PBMs of DGKκ (black) and η (orange) and different PDZ domains. **F**. Correlation between the binding affinities measured for DGKκ and η peptides. PDZ domains are ranked from the strongest to the lowest affinity from left to right for the first DGK (black) and the remaining non-common binders for the second DGK (orange) were ranked from right to left according to their affinities. R^2^: Pearson correlation coefficient, p: p-value of two-tailed t-test, red dotted lines: 95% confidence intervals, p*K*_d_ = −log_10_(*K*_d_ [M]).

The binding profiles of DGKι (type IV) and DGKη (type II) were also previously measured, albeit only using 133 PDZ domains. Interestingly, we found that DGKζ and DGKι share highly similar binding profile, as they bind almost the same proteins (common binders n=53, Jaccard similarity=0.83, Fig. 3C) with almost the same binding constants (Pearson correlation coefficient R^2^=0.88, two-tailed t-test p<0.0001, Fig. 3D). In the case of DGKη we observed low binding affinities (all measured p*K*_d_ values were below 5) just as in the case of DGKκ (Fig. 3E). No overlap between binders (common binders n=8, Jaccard similarity=0.2) or significant correlation between the affinities of the DGKκ and DGKη PDZ partners were identified (R^2^=0.30, two-tailed t-test p=0.16, Fig. 3F).

We compared the fragmentomic affinities of the DGKζ peptide from the ProfAff database and the affinities obtained for full-length proteins in the nHU experiment (Fig. S1 and 2D). Due to the quantitative nature of the nHU experiment, even affinity values below the binding threshold may be meaningful, but due to the low affinities we lose statistical power. For this reason, we included even values below the binding threshold in the case of nHU. We used estimated composite affinities from fragmentomic affinities for multi-PDZ domain-containing proteins (such as DLGs or SCRIB, see further details in Materials and Methods) *(25)*. The statistical correlation between the affinity values of the full-length proteins and the isolated domains was just below the threshold of α = 0.05, indicating a likely agreement (R^2^=0.50, p=0.052). Nonetheless, the lack of correlation can be attributed to the possibility that the full-length proteins in the native experiment may possess additional interaction sites that can influence binding, whereas isolated domains may behave differently when embedded in the full-length protein. Additionally, fragmentomic affinities were measured on individual domains that we combined in a simple manner to obtain a singular affinity for multi-PDZ proteins (see Materials and Methods), but in reality, the actual affinities may depend on the individual domain affinities in a more intricate manner.

Overall, for the DGKζ peptide, the nHU-MS allowed us to identify and quantify the interaction with 7 full-length partners. With AP-MS, we expanded the partner count to a total of 12, including 6 partners that were also detected in nHU-MS and an additional 6 partners. Our highly sensitive fragmentomic holdup approach revealed and quantified 55 partners for the DGKζ peptide and added 45 PDZ domain-containing proteins to the list of binders compared to the combined results of nHU-MS and AP-MS (Fig. 4).

**Figure 4.**
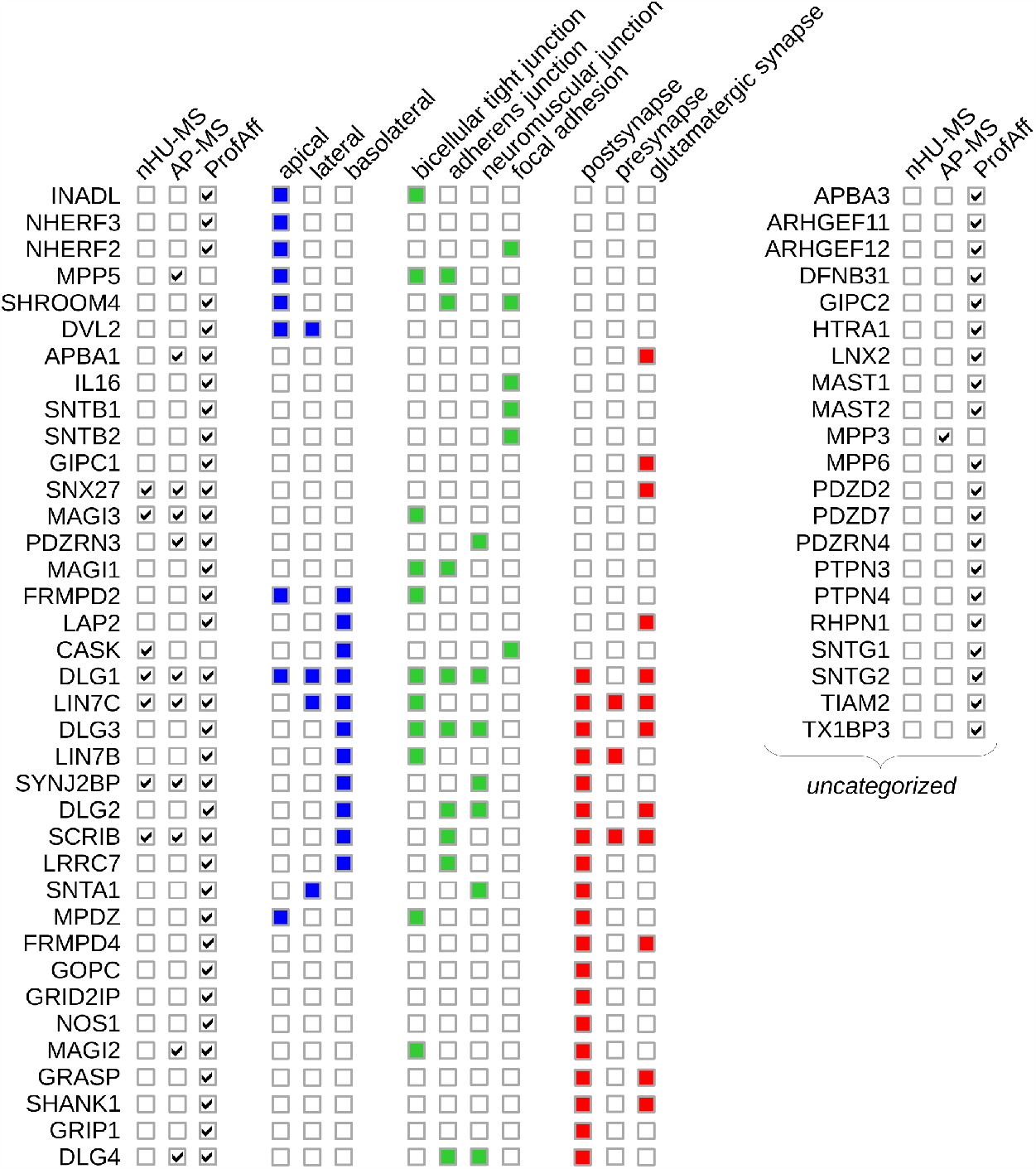
PDZ interaction partners of DGKζ are localized to different subcellular regions in epithelial and neuronal cells. Clustering based on their subcellular localizations (blue: epithelial membrane compartments, green: cell-cell and cell-matrix junctional regions, red: synaptic locations). Cellular compartment analysis of the binders of the DGKζ tail with DAVID tool with standard parameters.^33^

### Evolutionary conservation analysis of the PBM-like sequences in the DGK family

The PDZ domains are evolutionary highly conserved modules of the proteome, and their interactions can create significant selection pressure leading to evolutionarily conserved PBMs *(29)*. Therefore, we investigated the degree of conservation of PBMs among DGK proteins, which may indicate their binding ability to PDZ proteins. We collected DGK sequences across all the phylogenetic tree by sampling all major taxa equally and taking the Lifemap tool as a guide*(30)*. First, 176 sequences were collected from 39 species of different taxa using the Uniprot database. The domain architecture in the DGK family is highly conserved from multicellular organisms (as an example, we present here the domain structures of DGKs from human and brachiopod Lingula unguis, Fig. S2A and B, respectively), thus we used domain structure identification to separate the different types and enzymes in the first round using the NCBI conserved domain database *(31)*. Then, with careful manual selection and annotation, all DGKδ and alternative transcripts (mainly isoform 2 of DGKη) were removed. DGKδ contains a class II PDZ binding motif that seems highly variable among species, and the second isoform of DGKη does not contain the C-terminal PBM sequence. After selection, 114 sequences were kept from 37 species in total (54 type II and 60 type IV DGKs, Table S5). The classification was then validated by alignment and phylogram creation for all collected sequences, where the different types and enzymes were grouped together (Fig. S3). Note that the annotation of the enzymes is not complete and there might be species containing the whole set of enzymes, but we were unable to identify them in the database.

Human homologous DGKs appear very early on in the evolutional tree in the Eukarya superkingdom. Green plants (Viridiplantaea) have DGK proteins with similar enzyme mechanism but with different domain architectures and there are only three types of plant DGKs, all without PBMs *(32)*. Type IV DGKs seem to be the more ancient, a protein with the typical domain structure already present in some fungi (Table S5). The five types of DGKs with the typical domain structure can first be distinguished in early metazoans (representative species of the evolutional tree for type II and type IV DGKs in Fig. 5A and B, respectively). In Cnidaria there is already a type IV enzyme with a C-terminal PBM sequence closely resembling to the human DGKζ sequence (YETAV). The DGKζ and ι sequences can undoubtedly be identified in fish (Cypriniphysae). The PBM of the type IV enzyme in cartilaginous fish is identical to that in humans (QETAV). In contrast, the first type II enzyme can be identified in sponges (Porifera). From cartilaginous fish to marsupials, two types of type II DGK are already present that are similar to DGKι (e.g. they typically have a SAM domain). One type of these enzymes forms a well-separated group during alignments, which does not form a common group with either ι or δ, but is clearly transcribed from a separate gene. It is likely that these enzymes are closer to the ancient DGKκ and we therefore call them DGKκ-like enzymes. A DGKκ that is similar to human is firstly separable in Afrotheria. Human-like PBM sequences are identifiable in cartilaginous fish (TISEI, SMSEV), but they are highly variable among groups.

**Figure 5.**
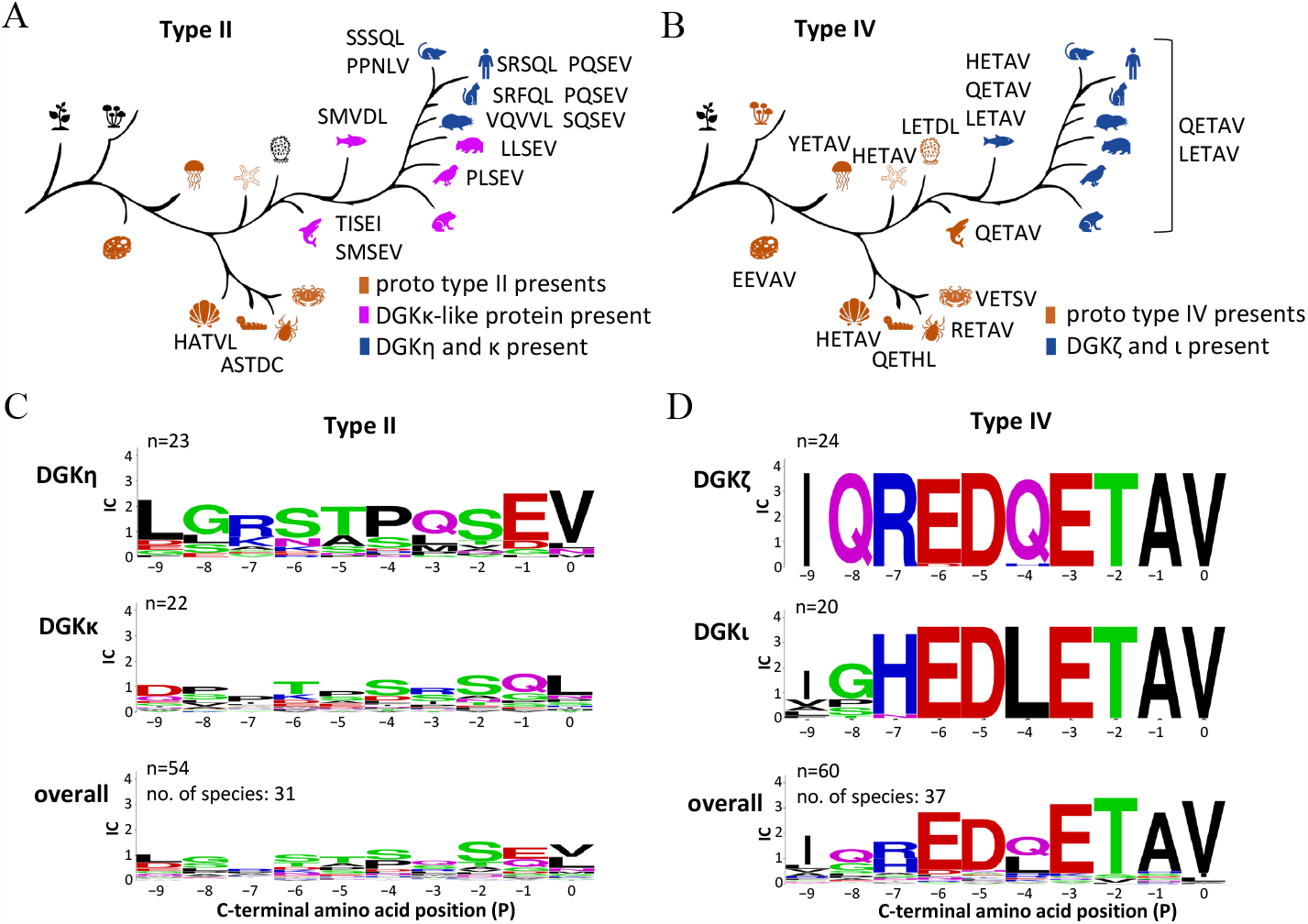
Evolutional conservation analysis of the PBMs among type II and type IV DGKs. **A-B.** The PBM of type II (A) and type IV (B) DGKs in representative species. DGK sequences were collected from all major clades from the evolutional tree and classified to different enzyme groups by using domain identification and phylograms based on the alignment of full sequences. Representative taxa and species from left to right: Viridiplantae (green plants), Fungi incertae sedis (fungi with unidentified origin), *Amphimedon queenslandica* (sponge), *Nematostella vectensis* (starlet sea anemone), *Lingula unguis* (lamp shell), *Caenorhabditis elegans* (roundworm), *Ixodes scapularis* (black-legged tick), *Daphnia magna* (waterflea), *Strongylocentrotus purpuratus* (purple sea urchin), *Ciona intestinalis* (sea vase), *Callorhinchus milii* (ghost shark), *Danio rerio* (zebrafish), *Xenopus tropicalis* (western clawed frog), *Taeniopygia guttata* (zebra finch), *Vombatus ursinus* (common wombat), *Chrysochloris asiatica* (cape golden mole), *Felis catus* (cat), *Mus musculus* (house mouse), *Homo sapiens* (human). **C-D**. Conservational sequence logos of PBMs in type II (C) and type IV (D) enzymes (created by WebLogo^34^).

We used the sequences collected to create a sequence logo in case of the different types and enzymes (Fig. 5C and D). Sequence logos indicate that the PBMs of type IV enzymes are highly conserved, while the PBMs of type II enzymes show weak conservation between different animal groups, validating functional differences that we have also observed with the combination of interactomic assays.

## Discussion

### Promiscuity and evolutional conservation of the PBMs in the DGK family

By using quantitative nHU-MS that is capable to determine equilibrium binding constants directly from cell extracts we determined the affinities of several PDZ domain-contacting full-length partners of DGKζ and ι from SH-SY5Y cell lysates. Among the significant partners of type II enzymes DGKκ, η and δ, we have not seen any PDZ domain-containing proteins, suggesting that these enzymes cannot be recruited by PDZ domains. We validated our findings using AP-MS on SH-SY5Y cell lysates where we did not find any notable partners for DGKκ, while we have identified several partners – most of them proteins containing the PDZ domain - for the DGKζ peptide. We also performed a careful meta-analysis of our recent high-throughput affinity survey, mapping the PDZ interactomic landscapes of PBMs in the DGK family. The ProfAff database include not only Boolean lists of interactions of the different DGK PBMs, but also precisely quantified site-specific affinity constants, which allow us to explore the binding preferences by ranking all potential target proteins across the proteome, as well as to precisely quantify the similarities or differences between the various members of the DGK family in multidimensional affinity spaces. Interestingly, both in the fragmentomic holdup (from the ProfAff database) and in nHU, the binding profiles of the two type IV DGKs, DGKζ and ι mediate highly similar affinities with PDZ domains. According to our results, type IV DGKs are highly promiscuous PDZ domain binders with many strong interactions, while type II DGKs do not have any strong binders in particular. We further validated our observations by evolutional conservation analysis, where we found that the PBMs of type II DGKs are not conserved across species, while type IV showed high conservation. Overall, we demonstrated that the PBMs of type II DGKs are rather nonfunctional or form low affinity complexes, while the binding profiles of the PBMs of type IV DGKs are highly similar, suggesting that in cells they may be the part of the same supramolecular complexes depending on the availability of the partner proteins.

### In-depth description of the PDZ-mediated interactions of DGKζ

Using a combination of approaches we described the most relevant partners of the C-terminal tail of DGKζ (Fig. 4). We have found that 58 PDZ proteins show detectable binding out of the 150 PDZ proteins encoded in the human genome. Not surprisingly, most of these partners were identified in the fragment-based holdup method (ProfAff), because this approach systematically probes interactions with all human PDZ domains and is significantly more sensitive than the nHU assay due to its more precise analytical approach *(25)*. However, not all PDZ proteins present in a given cell and therefore the complexes of DGKζ with PDZ proteins will depend on the cell type *(27)*. Since neuronal activities of DGKs are of high importance, we studied in depth the interactions of DGKζ in SH-SY5Y glioblastoma cell line.

With the nHU approach, we have determined several binding affinities with PDZ domain-containing proteins directly from SH-SY5Y cell extracts. Among them, the strongest affinities have been observed between SNX27, MAGI3, LIN7C, CASK and SYN2BP and the DGKζ peptide (Fig. 4). Although fragmentomic studies revealed high affinity interactions with the PDZ domains of syntrophins (SNTG1, SNTA1, SNTB1, SNTG2) and DLGs (DLG1, DLG4), these partners were either not detected in the nHU or did not show significant binding (DLG1, p*K*_app_=4.78). It is important to note that interactions below significance threshold can be meaningful in nHU, because fold change values in these experiments are physical constants and interactions with weak affinities will result in small fold changes that can be challenging to measure statistically significantly *(23)*. Consequently, due to the lack of enrichment of interaction partners, nHU could fail to identify those with abundances below the detection threshold of MS. To detect these binding partners from cell extracts, we used AP-MS, where enrichment of partner proteins aides their identification at the expense of the loss of affinity information provided by nHU. Overall, the combination of these different techniques gives a picture of the most important partners for the tail of DGKs in glioblastoma cell lines, but the actual complexes with the tail of DGKζ may differ in different cell types. For example, in neurons where synthrophins are highly expressed, these interactions may dominate complex formation with DGKs, but in other cell types the dominant partners may be other proteins such as SNX27, MAGI1 or DLG1.

Nevertheless, considering all putative partners of DGKζ – identified with any of our approaches – shows that it has a high preference to bind postsynaptic proteins, proteins found in glutamatergic synapse, proteins involved in cell-cell contacts, such as tight junctions or adherens junctions, that often also show strong polarization based on cellular compartment analysis (DAVID tool, Fig. 4) *(33)*. Given the biological importance of these features, the strong evolutionary conservation of the tails of type IV DGKs is not surprising and it marks a clear contrast with the type II DGKs who apparently lack their capacity to bind to the same targets, yet whose involvement in neuronal activities is equally important, but certainly not mediated by PDZ domains through their C-terminal tails.

## Materials and Methods

### Cell culturing and lysate preparation for AP-MS and nHU-MS

SH-SY5Y cells (ATCC, #CTR-2266) were cultured in RPMI 1640 medium without HEPES (Gibco, ref. 11872101) complemented with 10% FCS and 40 μg/mL gentamycin and kept in a 37 °C incubator, diluted twice a week 1/5.

For cell lysate preparation, 5 × 10^6^ cells were seeded onto a T-175 flask. When the cells reached confluence, they were placed on ice and washed once with sterile PBS. Then, 1 mL of ice-cold lysis buffer (50 mM Hepes-KOH pH 7.5, 150 mM NaCl, 1% Triton X-100, 1 × cOmplete EDTA-free protease inhibitor cocktail, 2 mM EDTA, 5 mM TCEP, 10% glycerol) was added to them and the cells were scraped from the surface of the flask and collected in 15 mL Falcon tubes. The lysate was sonicated 4 × 20 sec with 1 sec long pulses on ice and incubated on a roller mixer for 30 minutes at 4°C, then centrifuged at 12,000 g for 20 minutes at 4°C. The supernatant was kept and the total protein concentration was determined by standard Bradford method (Bio-Rad Protein Assay Dye Reagent #5000006) using a BSA calibration curve (MP BIomedicals #160069, diluted in lysis buffer) on a Bio-Rad SmartSpec 3000 spectrophotometer instrument. Lysates were diluted to 2 mg/ml and were snap-frozen in liquid nitrogen and stored at -80°C until the measurements.

### Peptide synthesis

HPLC purified (>95% purity) biotinylated 10-mer peptides of the five DGK proteins (DGKδ, DGKη, DGKκ, DGKζ, DGKι, gene names: DGKD, DGKH, DGKK, DGKZ, DGKI, respectively, Uniprot accession codes: Q16760-1, Q86XP1-1, Q5KSL6, Q13574-2, O75912, respectively) were chemically synthesized by an ABI 443A synthesizer with standard Fmoc strategy with the biotin group attached to the N-terminus via a trioxatridecan-succinamicacid (ttds) linker and were purified with high-performance liquid chromatography (LC, >95%purity). Predicted peptide masses were confirmed by MS. Peptide concentrations were determined based on their dry weight.

### Native holdup (nHU)

Native holdup with the peptide baits has been performed as described before *(23)*. Briefly, we saturated 25 μl streptavidin resin (Cytiva, Streptavidin Sepharose High Performance, ref. GE17-5113-01) with biotin (control) or peptides at 60 μM concentration in 6 resin volume holdup buffer containing 50 mM tris (pH 7.5), 300 mM NaCl, and 1 mM TCEP (0.22-μm filtered) for one hour at RT. According to our previous validation experiments, ∼10 μM bait concentration can be achieved by mixing 25 μl of this bait-saturated streptavidin resin with 100 μl of cell lysate *(23, 25)*. The remaining sites on the resin were fully depleted by adding 25 μl 1 mM biotin and incubated it for 10 min at RT. Resins were washed three times with 10 resin volume holdup buffer. Then, the saturated streptavidin resins were mixed with 100 μl of SH-SY5Y cell lysate (2 mg/ml) and incubated by gentle rolling for 2 hours at 4 °C. After the incubation, the resins were separated from the supernatants by brief centrifugation (15 s, 2,000 g) and the supernatants were kept for label-free quantitative MS analyses.

### Affinity purification (AP)

During the affinity purification experiment, 25 μl streptavidin resin was mixed with 165 μl biotin or biotinylated peptide at a concentration of 63 μM and incubated for 1 hour at room temperature in AP buffer (50 mM Tris pH8.5, 150 mM NaCl, 1 mM TCEP) to achieve resin saturation. The resins were washed once with 10x resin volume of AP buffer and depleted with 40 μl 1 mM biotin for 10 minutes at room temperature. Finally, resins were washed twice with 10 × resin volume of AP buffer. After centrifugation of the resins (800 rpm 2 min) the supernatant was completely removed and 1 mg (0.5 mL, 2 mg/ml) of cell lysate was added to each tube and incubated by gentle rolling for 2 hr at 4°C. The resins were then washed 3-times with 10x resin volume of Wash buffer (50 mM Tris pH8.5, 150 mM NaCl, 1 × cOmplete EDTA-free protease inhibitor cocktail, 2 mM EDTA, 1% Triton-X, 1 mM TCEP) and 3 times with AP buffer. After centrifugation (800 rpm 2 min) the supernatant was completely removed and the elution buffer (8 mM urea, 20 mM Tris pH8.5, 100 mM NaCl, 100 μM TCEP) was added at 3x resin volume and incubated for 30 min at room temperature mixing often. The samples were centrifuged at 2,000 g for 1 min, and the supernatant was kept. Then the elution step was repeated and the two supernatants were mixed and centrifuged again 2,000 g for 1 min to completely remove all the remaining resin. This final supernatant was kept for label-free quantitative MS analysis.

### Sample digestion for MS

The samples were precipitated with 20% TCA overnight at 4°C and centrifuged at 14,000 rpm for 10 minutes at 4°C. Protein pellets were washed twice with 1 mL of cold acetone and air dried. The protein extracts were solubilized in 8 M urea, reduced with 5 mM TCEP for 30 minutes, and alkylated with 10 mM iodoacetamide for 30 minutes in the dark. Double digestion was carried out at 37°C in 2 M urea with 1 μg endoproteinase Lys-C and 1 μg trypsin overnight. The peptide mixtures were then desalted on C18 spin-column and dried on Speed-Vacuum.

### LC-MS/MS analysis

Samples were analyzed using an Ultimate 3000 nano-RSLC (Thermo Scientific, San Jose California) coupled in line with LTQ-Orbitrap Elite (AP-MS) or Exploris (nHU-MS) mass spectrometers via nano-electrospray ionization source (Thermo Scientific, San Jose California). Peptide mixtures were injected in 0.1% TFA into a C18 Acclaim PepMap100 trap-column (75 μm ID × 2 cm, 3 μm, 100Å, Thermo Fisher Scientific) for 3 minutes at 5 μL/min with 2% ACN, 0.1% FA in H_2_O and then separated in a C18 Accucore nano-column (75 μm ID x 50 cm, 2.6 μm, 150Å, Thermo Fisher Scientific) at 220 nl/min and 38 °C with a 90 minutes linear gradient from 5% to 30% buffer B (A: 0.1% FA in H_2_O / B: 99% ACN, 0.1% FA in H_2_O), regeneration at 5% B. The mass spectrometer was operated in positive ionization mode, in data-dependent mode with survey scans from m/z 350-1500 acquired in the Orbitrap at a resolution of 120,000 at m/z 400. The 20 most intense peaks from survey scans were selected for further fragmentation in the Linear Ion Trap with an isolation window of 2.0 Da and fragmented by CID with normalized collision energy of 35% (TOP20CID method). Unassigned and single charged states were excluded from fragmentation. The Ion Target Value for the survey scans (in the Orbitrap) and the MS2 mode (in the Linear Ion Trap) were set to 1E6 and 5E3, respectively, and the maximum injection time was set to 100 ms for both scan modes. Dynamic exclusion was set to 20 s after one repeat count with mass width at ± 10 ppm. The raw LC tandem MS data have been deposited to the ProteomeXchange via the PRIDE database with identifier PXD042853.

### Data extraction from ProfAff database and analysis

For the comparative analysis of DGK PBM binding affinities we collected the data from the ProfAff database described previously (https://profaff.igbmc.science/) that contains equilibrium binding constants of PDZ-PBM interactions between purified PDZ domains and synthetic 10-mer C-terminal peptides *(25)*. To determine the Jaccard similarities, we used the built-in ‘compare’ function of the website. For profile and affinity comparisons the retrieved data can be found in Table S4. To determine correlation between the data sets we used linear regression fitting using only those affinity pairs where both motifs showed measurable affinity for the given PDZ domain in the database. The significance of the correlations was determined using two-sided Student’s t-test using the null hypothesis that the slope is 0 and the 95% confidence intervals were depicted using GraphPad Prism 7 software.

We calculated composite affinity values to estimate affinities for full-length proteins from domain affinities found in the ProfAff database in case of multi-PDZ domain-containing proteins as before *(25)*. This calculated value does not consider potentially important factors contributing to affinity, such as the formation of higher order multivalent complexes, but the lower limit of affinity can be estimated by the help of the formula *(25)*.

### Data analysis

For mass spectrometry data analysis, proteins were identified by database searching using SequestHT (Thermo Fisher Scientific) with Proteome Discoverer 2.4 software (PD2.4, Thermo Fisher Scientific) in the human fasta database downloaded from SwissProt. Precursor and fragment mass tolerances were set at 7 ppm and 0.6 Da respectively, and up to 2 missed cleavages were allowed. Oxidation (M, +15.995 Da) was set as a variable modification and Carbamidomethylation (C, + 57.021 Da) as fixed modification. Peptides and proteins were filtered with a false discovery rate (FDR) at 1%. Label-free quantification was based on the extracted ion chromatography intensity of the peptides. All samples were measured in technical triplicate. The measured XIC intensities were normalized on median intensities of the entire dataset to correct for minor loading differences. For statistical tests and enrichment calculations, not detectable intensity values were treated with an imputation method, where missing values were replaced by random values similar to 10% of the lowest intensity values present in the entire dataset. Unpaired two tailed t-test, assuming equal variance, were performed on obtained log_2_ XIC intensities. The hyperbolic binding threshold for nHU-MS was calculated as follows

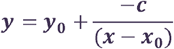

And for AP-MS as

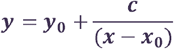

where *y* is the p-value threshold at the fold change of *x, c* is an empirically fixed curvature parameter at 1 for nHU-MS and 4 for AP-MS experiments, *y*_*0*_ is the minimal p-value threshold and *x*_*0*_ is the minimal fold change value for any given dataset. The minimal p-value was defined at 1.3−log_10_(p-value) and thus there is a >95% probability that all identified interaction partners are true interaction partners. The minimal fold-change value cutoff was set at 1 σ and was determined by measuring the width of the normal distribution of all measured fold changes in a given experiment. Note that this threshold can only be interpreted for interaction partners with fold change values of −x > x_0_ in the case of nHU-MS experiments or x > x0 in the case of AP-MS experiments. nHU-MS data are available in Table S1 and AP-MS data are available in Table S2.

For further statistical analyses and visualizations, GraphPad Prism 7 software was used. For domain structure and protein feature visualizations, ProFeatMap website tool was used *(28)*. Cellular compartment analysis was performed with the help of DAVID tool with standard parameters *(33)*.

### Collection of DGK sequences across the phylogenetic tree, classification and sequence logos

For the evolutionary conservation analysis of PBMs, DGK sequences were manually collected from all major clades of the phylogenetic tree using the Lifemap tool *(30)* as a guide and the Uniprot database. In total, 176 sequences were collected in the first round from 40 species. Careful manual selection was performed by removing all DGKδ and alternative transcripts without PBM. After the selection, 114 sequences were kept from which 54 identified as type II and 60 as type IV (Table S5). DGKs are classified into different types based on their domain structure, we used NCBI Conserved Domain Database Search *(31)* to identify the different types of DGKs and the different enzymes. For final validation, an alignment was performed using ClustalΩ multiple sequence alignment and a sequence phylogram was created using Jalview version 2.11.2.4 (Fig. S3). Sequence logos were created using WebLogo tool *(34)*. Note that DGKκ-like sequences were indicated as DGKκ during logo creation.

## Supporting information

Zambo_Supplementary_Materials

## Supplementary Materials

Fig. S1: Correlation between the measured apparent affinities in native holdup (nHU-MS) and the calculated composite affinities using the ProfAff database for the DGKζ peptide.

Fig. S2: The domain structure of DGKs is conserved among animals.

Fig. S3: The selected 114 DGK sequences used for evolutional conservation analysis of the PBMs.

Table S1: The results of native holdup mass spectrometry measurements with the C-terminal PBMs of DGKs.

Table S2: The results of affinity purification mass spectrometry measurements with the C-terminal PBMs of DGKζ and DGKκ.

Table S3: Comparision of p*K*_app_ (nHU-MS) and enrichment values (AP-MS) of significant partners for DGKζ.

Table S4: Comparision of PDZ-binding profiles of different DGKs using the ProfAff database. Table S5: DGK sequences used in evolutional analyses.

## Acknowledgement

We are grateful for the help of the cell culture platform at IGBMC in this project. We thank to Goran Bich for his kind help in figure generation using the ProFeatMap.

## Author’s contribution

BZ, GG, HM and GT conceptualized the study. BZ and GG designed and carried out the experiments and all the analyses. Peptide syntheses were performed by PE. MS experiments were performed by BM and LN. GT and HM obtained funding for the experiments. BZ wrote the original draft of the paper. All authors reviewed the manuscript.

## Data and materials availability

All data and materials needed to evaluate the conclusions in the paper are present in the paper and/or the Supplementary Materials and in public databases. Raw mass spectrometry data are available via ProteomeXchange with identifier PXD042853.

