## Supplementary material for "Comparative analysis of PDZ-binding motifs in the diacylglycerol kinase family": Zambo_Supplementary_Materials: SuppFig1.pdf

Supplementary Figure 1.

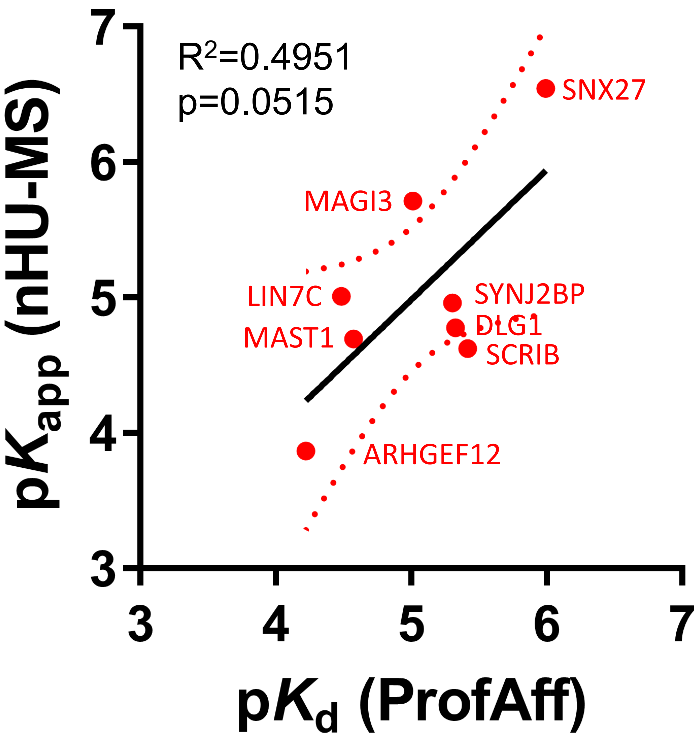


**Supplementary Figure 1. Correlation between the measured apparent affinities in native holdup (nHU-MS) and the calculated composite affinities using the ProfAff database for the DGKζ peptide.** $R^2$: Pearson correlation coefficient, p: p-value of two-tailed t-test, red dotted line: 95% confidence intervals.
