## Supplementary material for "Comparative analysis of PDZ-binding motifs in the diacylglycerol kinase family": Zambo_Supplementary_Materials: SuppFig2.pdf

Supplementary Figure 2.

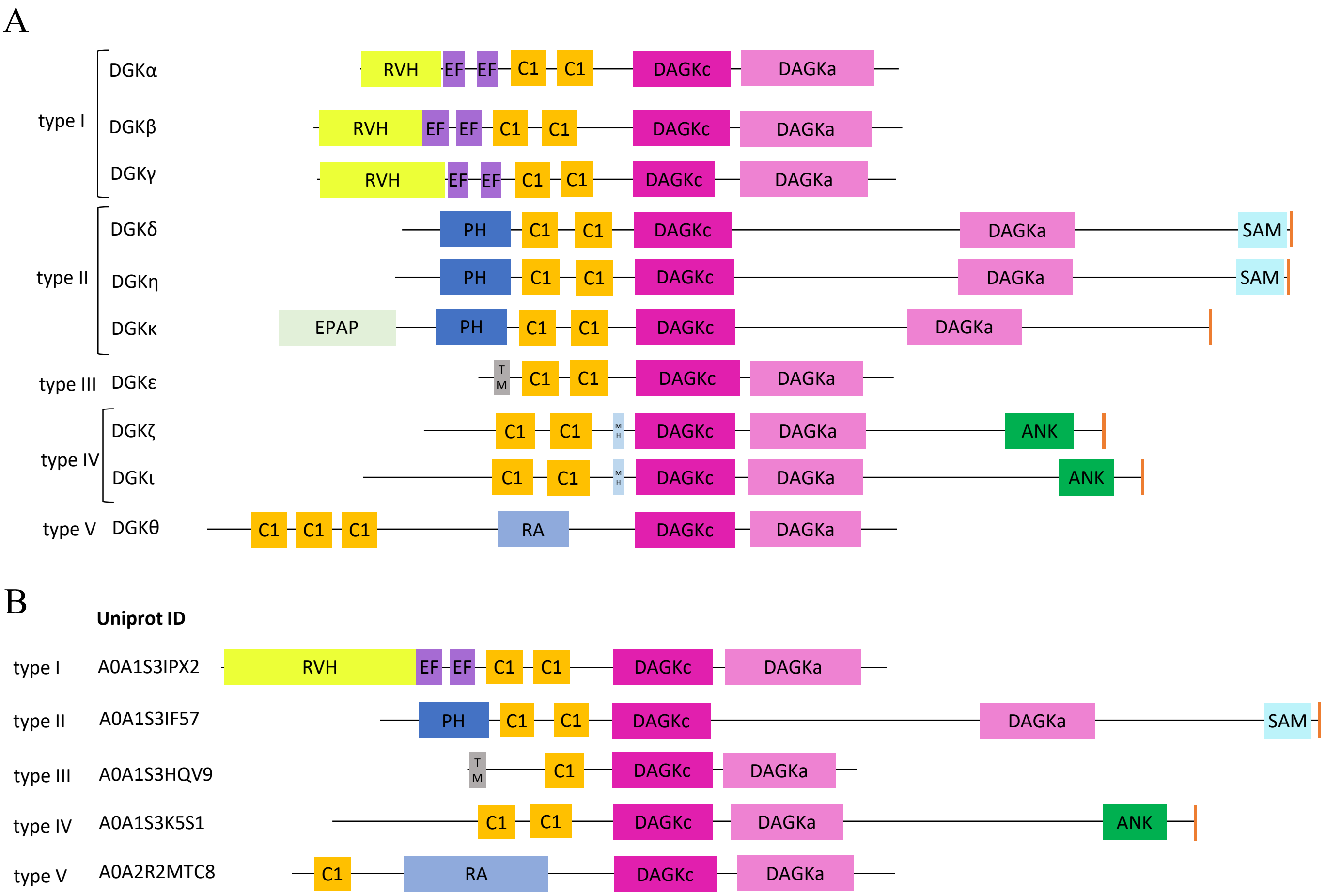


**Supplementary Figure 2. The domain structure of DGKs is conserved among animals. A.** The domain structure of human DGKs. **B.** The domain structure of brachiopod *Lingula unguis* DGKs. RVH: recovery homology domain, EF: $Ca^{2+}$-binding EF hand domain, C1: cysteine-rich C1 domain, DAGKc: diacylglycerol kinase catalytic domain, DAGKa: diacylglycerol kinase accessory domain, PH: pleckstrin homology domain, EPAP: E-P-A-P repeat region, SAM: sterile alpha motif, TM: transmembrane helix, MH: MARCKS homology domain, ANK: ankyrin repeat, RA: Ras association domain, orange C-terminal box: PBM.
