## Supplementary material for "Comparative analysis of PDZ-binding motifs in the diacylglycerol kinase family": Zambo_Supplementary_Materials: SuppFig3.pdf

Supplementary Figure 3.

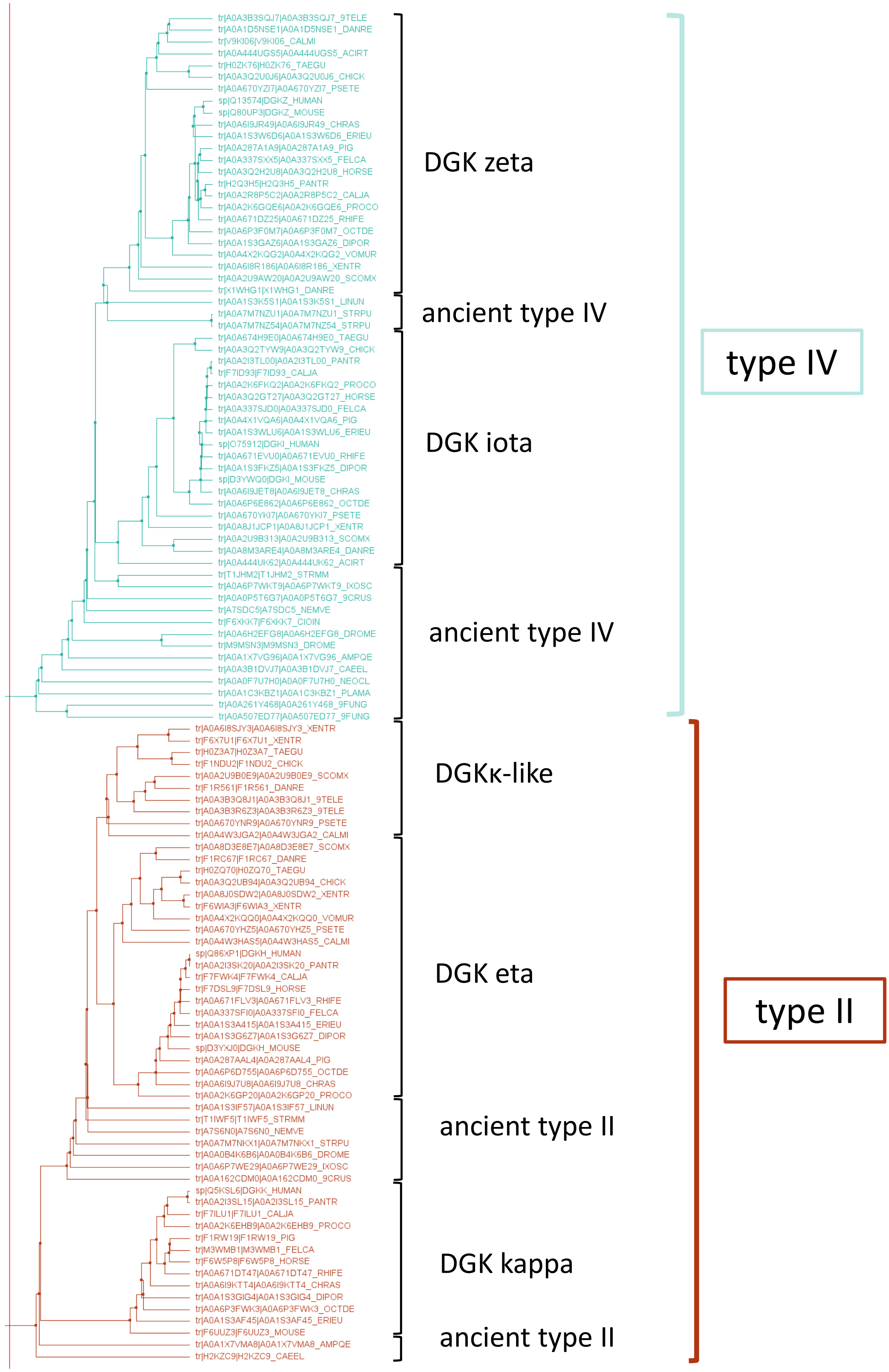


**Supplementary Figure 3. The selected 114 DGK sequences used for evolutional conservation analysis of the PBMs.** The phylogram was created by the alignment of the sequences using ClustalΩ and Jalview tools.
